# Mass cytometry identifies characteristic immune cell subsets in bronchoalveolar lavage fluid from interstitial lung diseases

**DOI:** 10.1101/2023.01.12.523734

**Authors:** Kentaro Hata, Toyoshi Yanagihara, Keisuke Matsubara, Kazufumi Kunimura, Kunihiro Suzuki, Kazuya Tsubouchi, Daisuke Eto, Hiroyuki Ando, Maki Uehara, Satoshi Ikegame, Yoshihiro Baba, Yoshinori Fukui, Isamu Okamoto

## Abstract

Immune cells have been implicated in interstitial lung diseases (ILDs), although their phenotypes and effector mechanisms remain poorly understood. To better understand these cells, we conducted an exploratory mass cytometry analysis of immune cell subsets in bronchoalveolar lavage fluid (BALF) from patients with idiopathic pulmonary fibrosis (IPF), connective-tissue disease (CTD)-related ILD, and sarcoidosis, using two panels including 64 markers. Among myeloid cells, we observed the expansion of CD14^+^ CD36^hi^ CD84^hi^ monocyte populations in IPF. These CD14^+^ CD36^hi^ CD84^hi^ subsets were also increased in ILDs with a progressive phenotype, particularly in a case of acute exacerbation (AEx) of IPF. Analysis of B cells revealed the presence of cells at various stages of differentiation in BALF, with a higher percentage of IgG memory B cells in CTD-ILDs and a trend toward more FCRL5^+^ B cells. These FCRL5^+^B cells were also present in the patient with AEx-IPF and sarcoidosis with advanced lung lesions. Among T cells, we found increased levels of IL-2R^+^TIGIT^+^ LAG3^+^ CD4^+^T cells in IPF, increased levels of CXCR3^+^ CD226^+^ CD4^+^ T cells in sarcoidosis, and increased levels of PD1^+^ TIGIT^+^ CD57^+^ CD8^+^ T cells in CTD-ILDs. Together, these findings underscore the diverse immunopathogenesis of ILDs.

## Introduction

Interstitial lung disease (ILD) is a broad term for a group of disorders characterized by varying degrees of inflammation and scarring or fibrosis of the lung (1). In the majority of cases, ILD is a chronic disease with a progressive scarring of lung tissue. Idiopathic pulmonary fibrosis (IPF) is the most common form of ILD and has a median survival rate of 3–5 years (2). Connective tissue disease (CTD), defined as systemic disorders characterized by autoimmune-mediated organ damage and circulating autoantibodies, is one of the common systemic diseases associated with ILD (3). Sarcoidosis is a granulomatous disorder of uncertain etiology, impacting various organ systems. Pulmonary involvement, including ILDs, represents the majority of morbidity and mortality associated with sarcoidosis (4).

It has been suggested that immune cells are involved in the pathogenesis of ILDs (4–7), with various subsets of immune cells potentially contributing to the development of ILDs, particularly macrophages and lymphoid cells. Macrophages, most abundant immune cells in the lungs(8), have been shown to play a key role in the initiation and progression of fibrotic responses(9). The ability of macrophages to alter their functional traits in response to external stimuli allows them to exhibit a range of biological impacts. Recent studies have showed that the existence of a unique population of macrophages in IPF lung explants (10)(11)(12) and murine models of pulmonary fibrosis (13)(14). T cells have also been implicated in the development of ILDs(6)(15). In addition, increasing evidence suggests that pathogenic B cells may contribute to the development of autoimmune diseases(16–18), although their role in the development of ILDs remains poorly understood. The inability to fully characterize immune cells in clinical specimens is partially due to a deficiency in available assays. While next generation sequencing analysis has facilitated the thorough analysis of cells, there is a constraint on the number that can be examined. It can be challenging to accurately classify the type of immune cells solely based on transcripts. Mass cytometry, on the other hand, offers a in-depth, high-dimensional description of the immune cell population through the simultaneous utilization of multiple markers (19), which will significantly alter our understanding of ILDs.

The objective of this study is to utilize mass cytometry to phenotype macrophages, B cells, and T cells in bronchoalveolar lavage fluid (BALF) cells from patients with IPF, CTD-ILD, and sarcoidosis. By employing both unbiased and manually classified methods, we functionally characterize subpopulations of myeloid cells, B cells, and T cells that are overexpressed in each disease, thereby identifying cells that may potentially affect disease progression.

## Materials and Methods

### Patients

Patients who underwent BALF collection and were newly diagnosed with IPF, CTD-ILD, and sarcoidosis between Jan 2017 and April 2022 at Kyushu University Hospital were eligible for enrollment in the study. The study was authorized by the Ethics Committee of Kyushu University Hospital (reference number 22117-00). The diagnostic criteria for IPF, CTD-ILD, and sarcoidosis were as described elsewhere (20–23)(24). Interstitial pneumonia with autoimmune features (IPAF) was included in CTD-ILD in this study. Disease progression was defined as follows: a relative decline of at least 10% in the predicted value of forced vital capacity (FVC), a relative decline of 5-10% in the predicted value of FVC accompanied by worsening respiratory symptoms or increased lung involvement on high-resolution CT imaging, or worsening respiratory symptoms and increased lung involvement within 24 months, as modified by the inclusion criteria of the INBUILD trial (25). Experimental and analytical workflow is shown in Figure 1.

**Figure 1.**
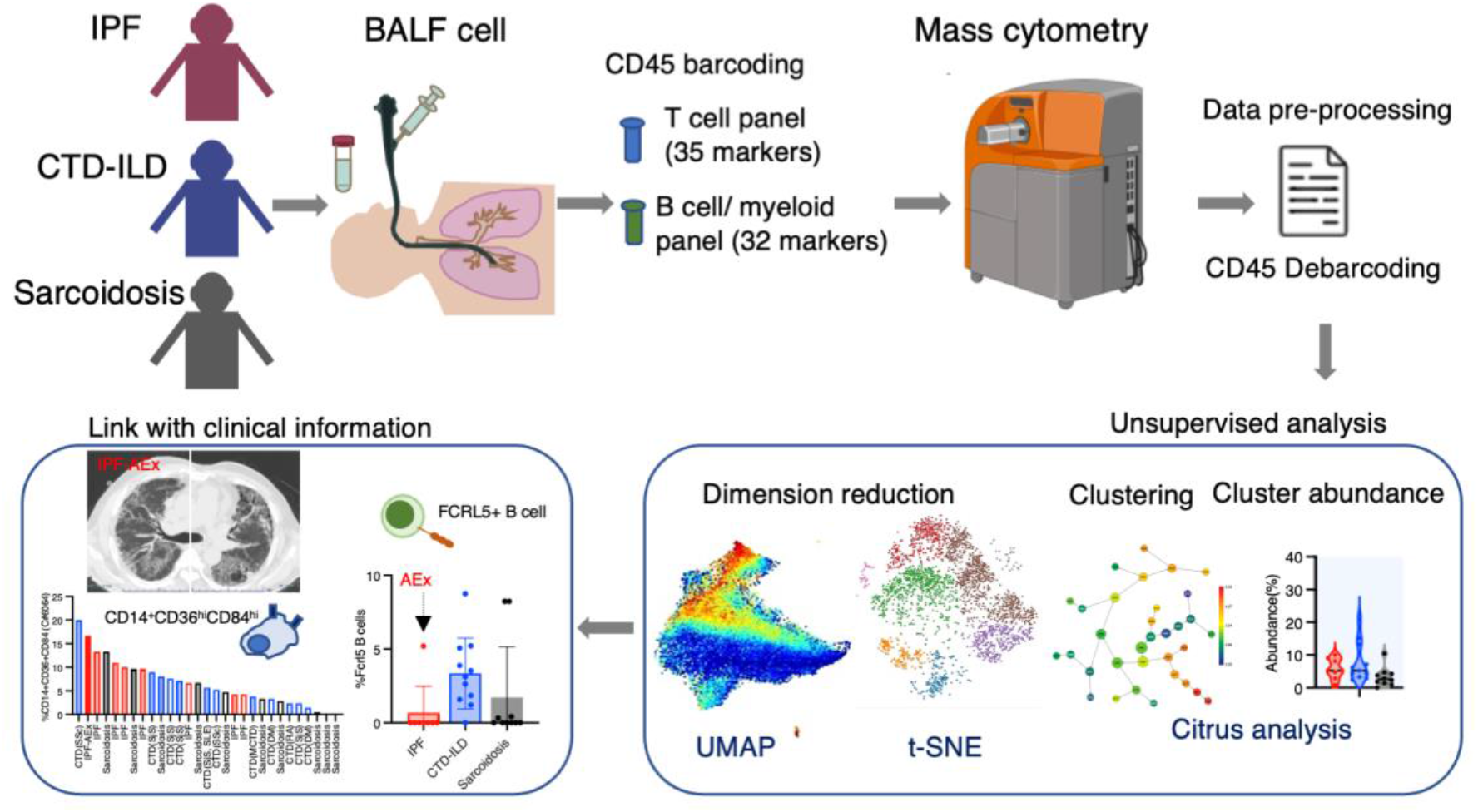
Graphical abstract of the study. Bronchoalveolar lavage fluid (BALF) samples were collected from patients with idiopathic pulmonary fibrosis (IPF), connective-tissue disease (CTD)-related ILD, and sarcoidosis. Following CD45 barcoding for individual sample identification, BALF cells were analysed with a T cell panel (35 markers) and B cell/myeloid cell panel (32 markers) using mass cytometry. The expansion of CD14^+^ CD36^hi^ CD84^hi^ monocyte populations were found in IPF and ILDs with a progressive phenotype. FCRL5^+^ B cells were increased in CTD-ILDs, acute exacerbation (AEx) of IPF, and sarcoidosis with advanced lung lesions.

### Mass cytometry

Antibodies were purchased either already metal-tagged (Standard Biotools) or in purified form (Supplementary Table 1). Purified antibodies were conjugated with metals using the Maxpar Antibody Labeling Kit (Standard Biotools) according to the manufacturer’s instructions and stored at 4°C. The cell labeling was performed as previously described (26). Briefly, cryopreserved BALF cells in Cellbanker1 (Takara #210409) were thawed in PBS and stained with Cell-ID^™^ Cisplatin-198Pt (Standard Biotools #201198, 1:2000 dilution) in PBS. Cells were incubated with FcR blocking reagent (Myltenyi, #130-059-901) and barcoded with each metal-labelled CD45 antibodies (Supplementary Table 1). After washing, CD45-labelled cells were mixed (maximum 6 samples together) and stained with APC-conjugated FCRL5 antibodies (Biolegend #340305)(for panel #2), followed by staining antibody cocktail (Panels #1 and #2, see Supplementary Table 1). The amount of antibodies were determined by preliminary experiments with metal minus one. After staining, cells were washed, fixed with 1.6% formaldehyde, and resuspended in Cell-ID Intercalator 103Rh (Standard Biotools #201103A) in Fix and Perm buffer (Standard Biotools) at 4°C overnight. For acquisition, cells were resuspended in MaxPar Cell Acquisition Solution (Standard Biotools #201240) containing one-fifth EQ Four Element Calibration Beads (Standard Biotools #201078) and were acquired at a rate of 200–300 events/second on a Helios mass cytometer (Standard Biotools). Files were converted to FCS, randomized, and normalized for EQ bead intensity using the Helios software. Concatenating fcs files in the same group into one file was conducted by FlowJo v10.8 (BD Biosciences). Manual gating, visualization of t-distributed stochastic neighbor embedding (viSNE) and Uniform manifold approximation and projection (UMAP) analysis, and Citrus analysis (27) were performed using Cytobank Premium (Cytobank Inc.).

### Data analysis

Live cells were selected by exclusion of cisplatin-positive cells and doublets. CD45^+^were selected and further analyzed. For myeloid cells, CD11b^+^CD11c^+^ cells were gated and Citrus algorithm was conducted with clustering channels of CD11b, CD11c, CD64, CD14, CD16, CD32, CD36, CD38, CD84, CD86, CD163, CD206, CD209, CD223, TIM-1, HLA-DR, CCR2, CCR5, ST2 using following parameters: association models = nearest shrunken centroid (PAMR), cluster characterization = abundance, minimum cluster size = 5%, cross validation folds = 5, false discovery rate = 1%. Two cases of CTD-ILDs (dermatomyositis-related, IgG4-related) were excluded due to low cell number to calculate clusters. UMAP analysis for myeloid cells was conducted with clustering channels of CD11b, CD11c, CD64, CD14, CD16, CD32, CD36, CD38, CD84, CD86, CD163, CD206, CD209, CD223, TIM-1, HLA-DR, CCR2, CCR5, ST2 using following parameters: numbers of neighbors = 15, minimum distance = 0.01. For B cells, CD3^-^CD64^-^ and CD19^+^ or CD138^+^ cells were gated and viSNE was conducted with clustering channels of CD19, CD38, CD11c, IgA, IgG, CD138, CD21, ST2, CXCR5, CD24, CD27, TIM-1, IgM, HLA-DR, IgD, FCRL5 on individual files and concatenated files using following parameters: iterations = 1000, perplexity = 30, theta = 0.5. Two cases of CTD-ILDs (dermatomyositis-related, IgG4-related) were excluded due to low cell number to calculate clusters. For T cells, CD2^+^CD3^+^ cells were gated on concatenated files and Citrus algorithm was conducted with clustering channels of CD4, CD5, CD7, CD8a, CD11a, CD16, CD27, CD28, CD44, CD45RA, CD45RO, CD49d, CD57, CD69, CD226, Fas, IL-2R, PD-L1, PD-L2, PD-1, OX40, TIGIT, TIM3, CTLA-4, CD223 (LAG-3), BTLA, ICOS, ST2, CCR7, CXCR3, HLA-DR using following parameters: association models = nearest shrunken centroid (PAMR), cluster characterization = abundance, minimum cluster size = 5%, cross validation folds = 5, false discovery rate = 1%. viSNE for T cells was conducted with clustering channels of CD4, CD5, CD7, CD8a, CD11a, CD16, CD27, CD28, CD44, CD45RA, CD45RO, CD49d, CD57, CD69, CD226, Fas, IL-2R, PD-L1, PD-L2, PD-1, OX40, TIGIT, TIM3, CTLA-4, CD223 (LAG-3), BTLA, ICOS, ST2, CCR7, CXCR3, HLA-DR using following parameters: iterations = 1000, perplexity = 30, theta = 0.5.

### Statistical analysis

Statistical analysis was conducted using one-way analysis of variance (ANOVA) followed by Dunnett’s multiple comparison test with the use of GraphPad Prism 8. A P-value <0.05 was considered statistically significant. The statistical parameters for Citrus argolism are described in the *Data Analysis* section.

## Results

### Patient characteristics

We analyzed 8 cases of IPF, 13 of CTD-ILD, and 10 of sarcoidosis (Table 1). Among the IPF cases, one patient experienced acute exacerbation (AEx) of IPF on admission. Thirteen cases of CTD-ILDs were observed, comprising Sjogren’s syndrome (n = 3), dermatomyositis (n = 3), systemic sclerosis (n = 2), mixed connective tissue disease (n = 1), systemic lupus erythematosus complicated with adult-onset Still’s disease (n = 1), IgG4-related disease (n = 1), rheumatoid arthritis (n = 1), and IPAF (n = 1). Differential cell counts for BALF revealed lymphocytosis in 1 out of 8 cases (12.5%) of IPF as well as in 7 out of 13 cases (53.8%) of CTD-ILD and 7 out of 11 cases (63.6%) of sarcoidosis when the cut-off for the percentage of lymphocytes was set to >20%.

**Table 1.** Characteristics of the study population. Data for age and BALF analysis are presented as mean±SD. IPF: idiopathic pulmonary fibrosis; CTD-ILD: connective tissue disease-related ILD; BALF: bronchoalveolar lavage fluid.

|  | IPF | CTD-ILD | Sarcoidosis |
| --- | --- | --- | --- |
| Number | 8 | 13 | 10 |
| Age | 72.0±7.6 | 66.2±15.1 | 48.7±15.2 |
| Male % (n) | 100 (8) | 38.5 (5) | 60 (6) |
| BALF cell differentiation (%) |  |  |  |
| Macrophage | 82.2±10.5 | 65.1±25.7 | 55.6±29.9 |
| Neutrophil | 6.2±3.9 | 7.5±18.1 | 1.9±2.7 |
| Lymphocyte | 9.5±9.4 | 26.0±23.2 | 42.0±31.0 |
| Eosinophil | 2.0±1.5 | 1.4±1.5 | 0.6±0.5 |
| CD4/CD8 ratio | 4.1±2.6 | 4.4±6.7 | 9.6±10.6 |

### Expansion of CD14^+^CD36^hi^CD84^hi^ monocytes in BALF from patients with IPF

To investigate myeloid cell populations (identified as CD45^+^CD11b^+^CD11c^+^) that could provide insight into ILD conditions, we utilized the Citrus algorithm to distinguish differently abundant myeloid cell subpopulations in IPF, CTD-ILD, and sarcoidosis through the analysis of 19 parameters (Figure 2A, all parameters can be seen in Supplementary Figure 1). Our analysis identified 33 clusters of myeloid cells, of which 23 were significantly differentiated between the groups (Supplementary Figure 1). Clusters #6074, #6025, #6066, and #6054 were prevalent in sarcoidosis and characterized by CD64^+^ CD11b^lo^ CD14^-^ CD223(LAG3)^+^ HLA-DR^+^ CD163^hi^ expression (Figure 2B). Cluster #6059, which was abundant in CTD-ILD, was marked by CD64^+^ CD11c^+^ CD11b^+^ CD38^hi^ expression (Figure 2B). Clusters #6075, #6064, and #6056, which were prevalent in IPF, were comprised of CD64^+^ CD11b^hi^ CD11c^hi^ CD14^+^ CD36^hi^ CD84^hi^ monocyte subpopulations (Figure 2B). A recent report indicated that CD36^hi^ CD84^hi^ macrophages were increased in IPF compared to control and COPD lungs (28), which is consistent with our findings. Notably, the CD14^+^ CD36^hi^ CD84^hi^ monocyte subpopulation was also increased in ILDs with a progressive phenotype (Figure 2D), suggesting that these cells may have pathogenic properties.

**Figure 2.**
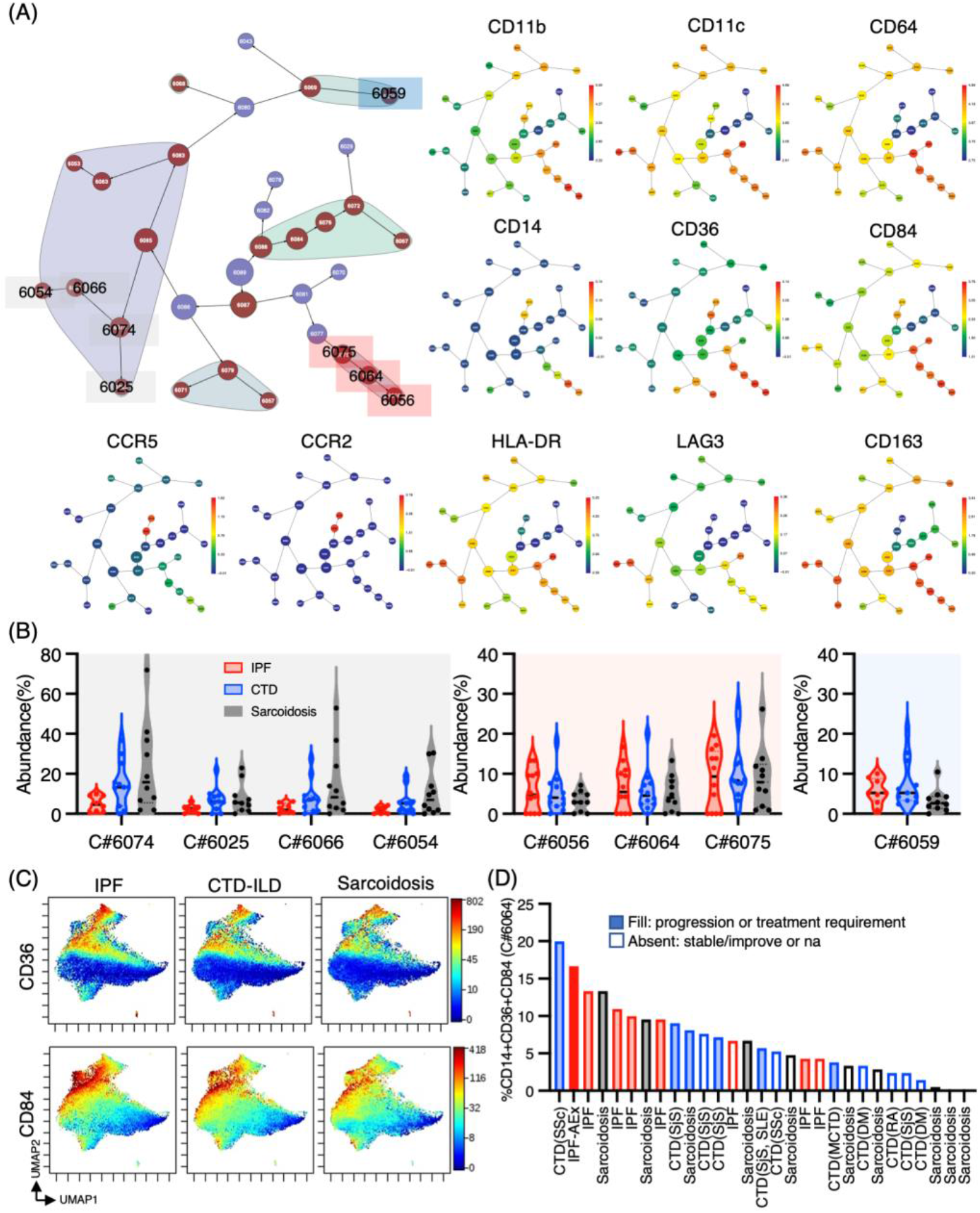
Characterization of myeloid cell subsets in BALF from patients with IPF, CTD-ILD, and sarcoidosis. (A) Citrus network tree visualizing the hierarchical relationship and intensity of each markers between identified myeloid cell populations gated by CD45^+^CD11b^+^ CD11c^+^ from IPF (n = 8), CTD-ILD (n = 11), and sarcoidosis (n = 10). Circle size reflects number of cells within a given cluster. (B) Citrus-generated violin plots for eight representative and differentially regulated populations. Each cluster number (C#) corresponds to the number shown in panel (A). All differences in abundance were significant at false discovery rate < 0.01. (C) Uniform manifold approximation and projection (UMAP) of myeloid cells showing cell distributions and CD36 or CD84 expression. (D) Correlation of cluster #6064 (CD14^+^ CD36^hi^ CD84^hi^) abundance in myeloid cell populations in individual samples. Information for disease with clinical progression is also shown. IPF: idiopathic pulmonary fibrosis, CTD-ILD: connective-tissue disease-related interstitial lung disease, AEx: acute exacerbation, SSc: systemic sclerosis, SjS: Sjogren syndrome, SLE: systemic lupus erythematosus, RA: rheumatoid arthritis, DM: dermatomyositis, na: not accessed.

### B cell subpopulations in the lungs of ILDs

Next, we sought to determine whether there were differential representations of B cell subsets in BALF cells of individuals with ILDs. Utilizing CD45^+^CD3^-^CD64^-^ and CD19^+^ or CD138^+^ as gating parameters, we were able to identify both B cells and plasma cells. It was observed that B cells tended to be more prevalent in individuals with CTD-ILDs and sarcoidosis compared to those with IPF, although the percentages of B cells/plasma cells remained low across all groups (Figure 3A). A tSNE analysis of 17 parameters among B cells/plasma cells revealed the presence of various B cell subpopulations, including IgD^+^ naïve B cells, IgM^+^ memory B cells, IgG^+^ memory B cells, IgA^+^ memory B cells, IgD^-^CD27^-^ double negative (DN) B cells, plasmablasts, and plasma cells (Figure 3B). This marked the first time that these B cell subpopulations were identified. The abundance of these subpopulations varied between groups, with particularly high levels of IgG memory B cells observed in individuals with CTD-ILDs compared to the other groups (Figure 3C).

**Figure 3.**
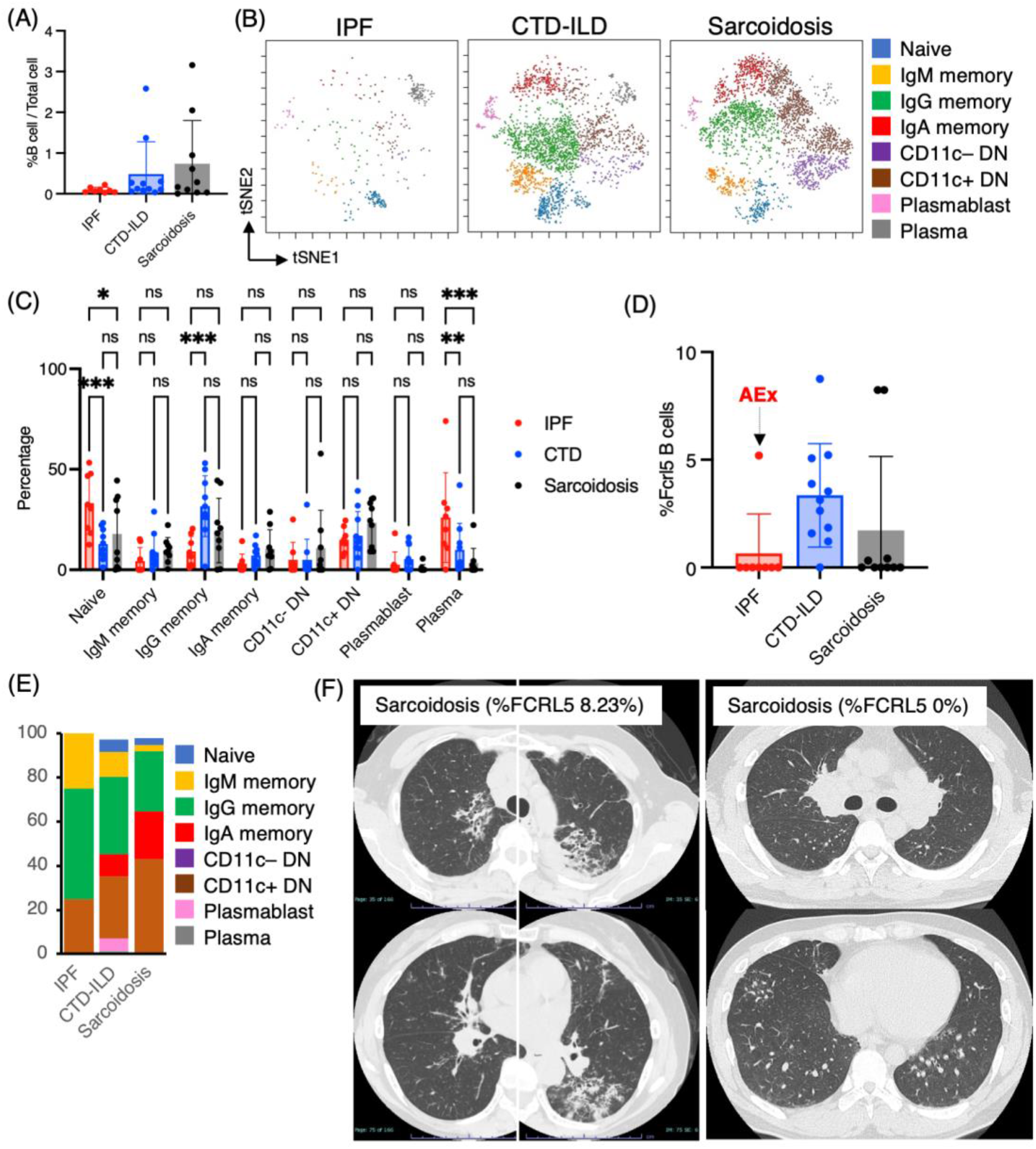
Characterization of B cell subsets in BALF from patients with IPF, CTD-ILD, and sarcoidosis. (A) Percentage of B cells and plasma cells in CD45^+^ BALF cells. (B) t-stochastic neighborhood embedding (t-SNE) plots of concatenated samples visualizing distribution of B cell sub-populations in CD64^-^CD3^-^ and CD19^+^ or CD138^+^ gated B cells in BALF from patients with IPF, CTD-ILD, sarcoidosis. Naive B cells are defined by CD19^+^IgD^+^, IgM memory B cells: CD19^+^ IgM^+^ CD27^+^, IgG memory B cells: CD19^+^IgG^+^ CD27^+^, IgA memory B cells: CD19^+^ IgA^+^ CD27^+^, CD11c^-^ double negative (DN) B cells: CD19^+^ CD11c^-^ IgD^-^ CD27^-^, CD11c^+^ DN B cells: CD19^+^ CD11c^+^ IgD^-^CD27^-^, plasmablasts: CD19^+^ CD27^+^ CD38^+^ CD138^-^, plasma cells: CD19^-^ CD138^+^ and IgG^+^ or IgA^+^. (C) Percentage of B cell sub-populations in IPF, CTD-ILD, and sarcoidosis. Graphical plots represent individual samples. (D) Percentage of FCRL5 expressing B cells in total B cells. (E) Proportion of each B cell subsets in FCRL5 expressing B cells. (F) Representative chest CT images of patients with sarcoidosis exhibiting a high percentage of FCRL5 B cells and a low percentage of FCRL5 B cells in BALF.

Recent evidence indicated that CD11c^+^ DN B cells and FCRL5^+^ B cells are involved in the development of autoimmune conditions(16)(17)(18)(29). In this study, we therefore sought to examine these B cell subsets. While we did not observe any differences in CD11c^+^ DN B cells between individuals with IPF, CTD-ILDs, and sarcoidosis (Figure 2C), there was a tendency for FCRL5^+^ B cells to increase in CTD-ILDs relative to the other groups (Figure 3D). FCRL5^+^ B cells were mostly IgG^+^ or IgA^+^ in each group (IPF: 75%, CTD-ILD: 80.3%, Sarcoidosis: 91.7%)(Figure 3E), suggesting that these FCRL5^+^ B cells were class-swithced B cells. Interestingly, there were 0% of FCRL5^+^ B cells in all IPF cases except for one, which was AEx of IPF. The higher FCRL5^+^ B cell percentage in sarcoidosis was associated with progressive lung involvement (Figure 3F). Conversely, a patient with sarcoidosis exhibiting a low percentage of FCRL5^+^ B cells, bilateral hilar lymphadenopathy, and small nodules on CT (Figure 3F) demonstrated marked improvement on follow-up. These findings suggest that FCRL5^+^ B cells may possess pathogenic properties in CTD-ILDs as well as in the context of AEx of IPF and sarcoidosis.

### Characteristic T cell subpopulations in the lungs of ILDs

In addition to myeloid cell and B cell subsets, we investigated whether subsets of T cells were differentially existed in ILDs. We utilized the Citrus algorithm to distinguish differently abundant T cell (defined as CD45^+^ CD2^+^ CD3^+^) subpopulations in IPF, CTD-ILD, and sarcoidosis through the analysis of 31 parameters (Figure 4, Supplementary Figure 4). Our analysis identified 31 clusters of T cells, of which 9 were significantly differentiated between the groups. Cluster #5508 was prevalent in sarcoidosis and characterized by CD4^+^ CD226^+^ CXCR3^+^. Cluster #5520, which was prevalent in IPF, was comprised of CD4^+^ IL-2R^+^ TIGIT^+^ LAG3^+^, considered as CD4^+^regulatory T cells (Tregs) (6). Cluster #5527, which was abundant in CTD-ILDs, was marked by CD8^+^ CD57^+^ PD-1^+^ TIGIT^+^.

**Figure 4.**
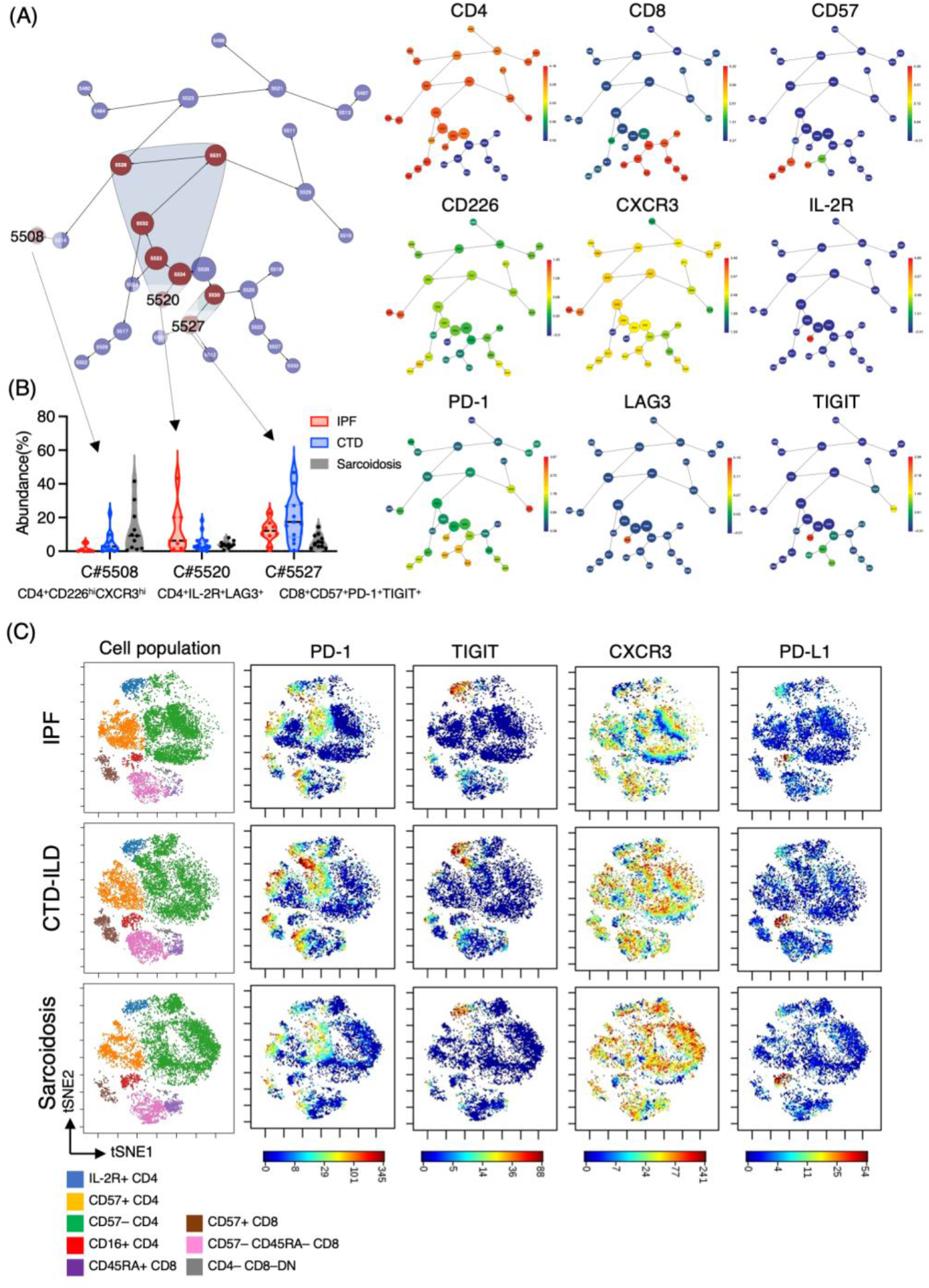
Characterization of T cell subsets in BALF from patients with IPF, CTD-ILD, and sarcoidosis. (A) Citrus network tree visualizing the hierarchical relationship and intensity of each markers between identified T cell populations gated by CD45^+^CD2^+^ CD3^+^ in BALF from IPF (n = 8), CTD-ILD (n = 13), and sarcoidosis (n = 10). Circle size reflects number of cells within a given cluster. (B) Citrus-generated violin plots for three representative and differentially regulated populations. Each cluster number (C#) corresponds to the number shown in panel (A). All differences in abundance are significant at false discovery rate < 0.01. (C) t-stochastic neighborhood embedding (t-SNE) plots of concatenated samples visualizing distribution of T cell sub-populations and PD-1, TIGIT, CXCR3, and PD-L1 expression.

### Immunological phenotypes in a patient with acute exacerbation of IPF

One of the patients suffering from IPF experienced an acute exacerbation. The patient complained of worsening dyspnea upon exertion and a dry cough. Upon admission, chest CT images revealed the emergence of bilateral diffuse ground glass opacities superimposed on a honeycomb pattern, accompanied by peripheral traction bronchiectasis primarily in the basal lungs (Figure 5A). BAL was conducted for differential diagnosis. Analysis of BALF revealed no evidence of infection. BALF cell differentiation showed a preponderance of macrophages (Figure 5B). The patient was diagnosed with an acute exacerbation of IPF and treated with pulse methylprednisolone, tacrolimus, antibiotics, and recombinant thrombomodulin. Upon admission, he required 5 L/min of oxygenation at rest. He showed improvement, requiring only 0.5 L/min of oxygenation, and was discharged three weeks after admission. We compared mass cytometry analysis of cell subpopulations between AEx (n = 1) and other cases (n = 7) of IPF. The proportion of CD14^+^ CD36^hi^ CD84^hi^ monocyte populations (clusters #6056, #6064, #6075) in myeloid cells was highest in AEx compared to stable conditions in IPF (Figure 5C). A UMAP of myeloid cells showed increased CD36 and CD84 expression in AEx of IPF (Figure 5D). On the contrary, the proportions of myeloid clusters #6025 and #6054, characterized by CD64^+^ CD11b^lo^ CD14^-^ CD223 (LAG3)^+^HLA-DR^+^ CD163^hi^ expression were lowest in AEx of IPF. t-SNE analysis of T cells showed a decreased proportion of CD8^+^ T cells and an increased proportion of CD4^+^CD57^-^ CD7^+^ CD44^+^ PD1^-^ subsets in AEx of IPF (Figure 4E).

**Figure 5.**
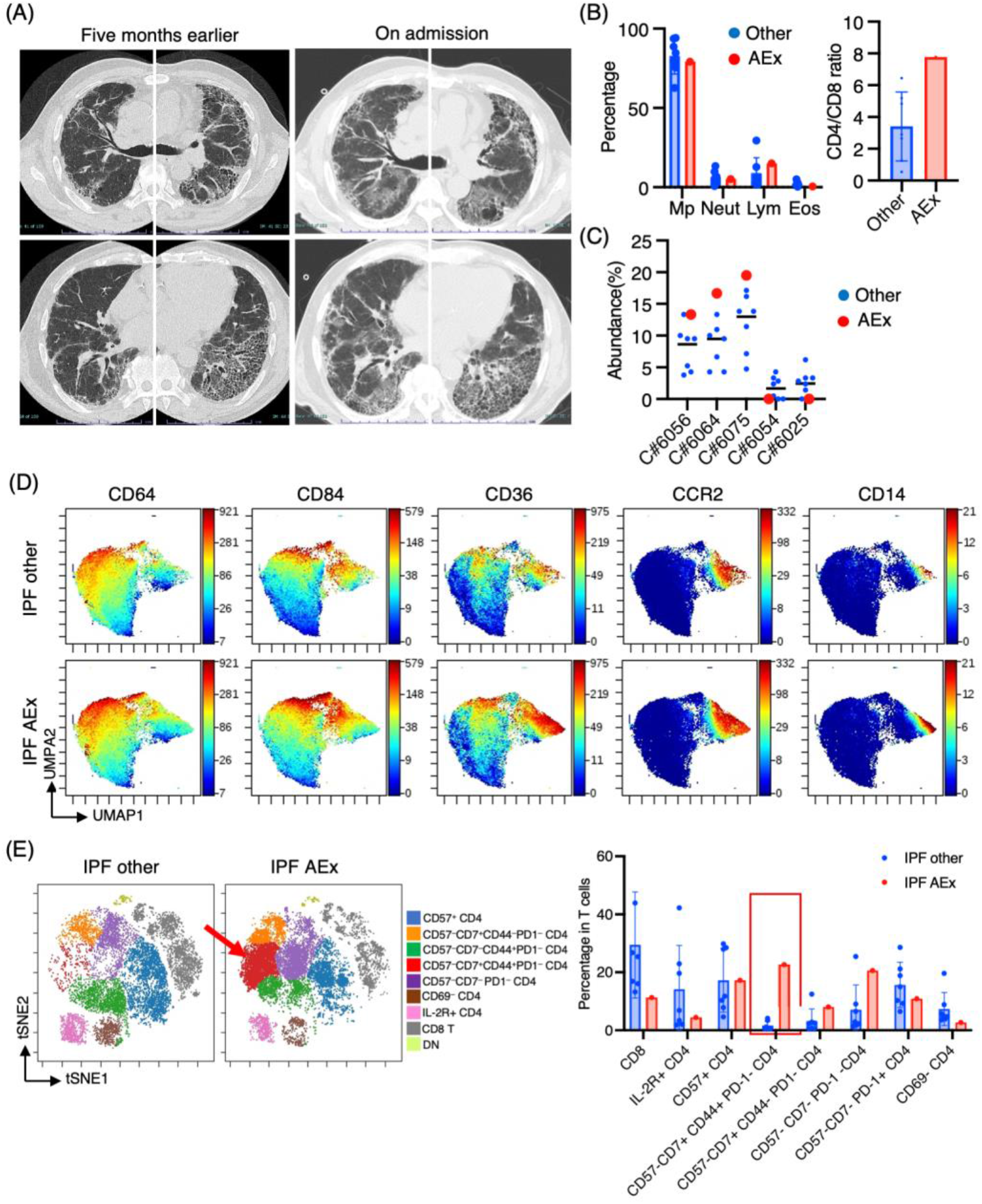
Immunological phenotypes in a patient with acute exacerbation of IPF. (A) Chest computed tomography images of the patient upon admission reveal the emergence of bilateral diffuse ground glass opacities superimposed on a honeycomb pattern, accompanied by peripheral traction bronchiectasis primarily in the basal lungs. (B) Comparison of BALF cell differentiation and CD4/CD8 ratio between patients with a patient experiencing an acute exacerbation (AEx) of the condition and the other cases of IPF. (C) Citrus-generated plots for myeloid sub-populations in IPF patients with stable condition and AEx. Each cluster number (C#) corresponds to the number shown in Figure 1A. (D) A uniform manifold approximation and projection (UMAP) of myeloid cells (CD45^+^CD11b^+^CD11c^+^ gated) showing cell distributions and each marker expression in BALF cells from concatenated samples with AEx and other cases of IPF. (E) A t-stochastic neighborhood embedding (t-SNE) plots visualizing distribution of T cell sub-populations in BALF T cells (gated as CD45^+^CD2^+^CD3^+^) from patients with AEx and other cases of IPF. Double negative (DN) T cells were defined as CD4^-^CD8^-^T cells.

## Discussion

We have here demonstrated the characteristic immune cell subpopulations present in BALF from patients with IPF, CTD-ILDs, and sarcoidosis. Our analysis revealed an expansion of CD14^+^CD36^hi^CD84^hi^ monocytes in patients with IPF, an increase in FCRL5^+^ B cell in patients with CTD-ILDs and AEx of IPF, increased levels of IL-2R^+^TIGIT^+^ LAG3^+^ CD4^+^ T cells in IPF, increased levels of CXCR3^+^ CD226^+^ CD4^+^ T cells in sarcoidosis, and increased levels of PD1^+^ TIGIT^+^ CD57^+^ CD8^+^ T cells in CTD-ILDs.

CD36 is a scavenger receptor expressed on the surface of immune and non-immune cells that acts as a signaling receptor for damage-associated molecular patterns (DAMPs) and pathogen-associated molecular patterns (PAMPs) and also serves as a transporter for long chain free fatty acids (30). CD84 is an immunoreceptor expressed on the surface of various immune cells that regulates a range of immunological processes, including T cell cytokine secretion, natural killer cell cytotoxicity, monocyte activation, autophagy, T–B interactions, and B cell tolerance at the germinal center checkpoint (31). Ayaub *et al*. recently reported that CD36^hi^ CD84^hi^ macrophages were expanded as a specific subpopulation of macrophages in the lungs of patients with IPF compared to healthy or chronic obstructive pulmonary disease lungs using single cell RNA-sequencing, mass cytometry, and flow cytometry (28). Ayaub *et al*. demonstrated that these CD36^hi^ CD84^hi^ macrophages expressed both alveolar and interstitial lung macrophage markers (HLA-DR^+^, CD11b^+^, CD206^+^), which is consistent with our results. Importantly, our study demonstrated these macrophage populations can be detected from BALF samples, which is less invasive compared to a lung biopsy and more suitable for clinical applications. We further determined that these CD36^hi^ CD84^hi^ macrophages were CD14 positive, indicating that these cells originated from monocytes.

We discovered these CD36^hi^ CD84^hi^ monocyte subpopulations were highest in AEx of IPF among all IPF cases. Nakashima *et al*. demonstrated that pulmonary fibrosis led to significant alterations in the bone marrow, including the expansion and activation of monocytic cells, which enhanced fibrosis upon subsequent lung injury (32). The pathobiology of IPF AEx is likely triggered by an acute event that leads to widespread acute lung injury, along with the acceleration of underlying chronic factors contributing to the fibrotic process (23). From these evidences, we speculate that these CD36^hi^ CD84^hi^ monocyte subpopulations may be involved in accelating AEx of IPF. Interestingly, these CD36^hi^CD84^hi^ CD14^+^ monocyte populations were ST2^+^ and CCR5^+^but distinct from CCR2^+^ monocyte populations (Figure 1A). Liang *et al.* demonstrated that the expression of CCL2, a ligand for CCR2, suppressed bleomycin-induced pulmonary fibrosis in mice (33). In addition, it has been shown that CCR2 deficiency in a mouse model of silica-induced pulmonary fibrosis resulted in an expansion of the fibrotic area (34), suggesting that CCR2^hi^ monocyte-derived macrophages may have a suppressive role in fibrosis. Increased monocyte count in blood samples has been identified as a cellular biomarker for poor outcomes in fibrotic diseases, including IPF (35). A more detailed analysis of subsets of monocytes in the blood may be more useful for predicting the prognosis of ILDs.

FCRL5, encoded by the immunoglobulin superfamily receptor translocation-associated 2 *(IRTA2)* gene, is a member of the Fc receptor-like family and its expression is mainly restricted to B cells. It has nine extracellular immunoglobulin domains and two immunoreceptor tyrosine-based inhibitory motifs as well as one presumed immunoreceptor tyrosine-based activation motif in its cytoplasmic tail (36). FCRL5 signaling, in conjunction with B cell receptor activation and TLR9 engagement, can lead to B-cell proliferation, activation, isotype switching, and the production of IgG- and IgA-positive B cells (29). Higher FCRL5 expression was shown to predict response to rituximab in rheumatoid arthritis (37), and granulomatosis with polyangiitis and microscopic polyangiitis (16). There are reports that single nucleotide variants in the FCRL5 gene increase an individual’s predisposition to multiple sclerosis (38) or SLE (39). These evidences indicate the pathogenic role of FCRL5 on autoimmunity. In this study, we have demonstrated increased FCRL5^+^ B cells in CTD-ILDs and a case of AEx of IPF. Recent study has demonstrated that rituximab was not inferior to cyclophosphamide to treat patients with CTD-ILDs with fewer adverse events (40). Collectively, higher FCRL5^+^ B cells in BALF could be used as a biomarker as a rationale for using rituximab.

We observed higher levels of IL-2R^+^ CD4^+^ T cells in patients with IPF compared to those with CTD-ILDs and sarcoidosis. These IL-2R^+^ CD4^+^ T cells likely represent Tregs, given that IL-2R^+^ expressing CD4+ T cells are a sole, distinct T cell subpopulation characterized by highest FOXP3 expression between T cell subsets (6). In the study by Serezani *et al*.(6), the proportions of CD4 Tregs were higher tendency in IPF compared to controls. Increased activated Treg proportion was correlated with the severity of IPF (41). However, the precise roles of Tregs in the development of pulmonary fibrosis have not been fully elucidated, and the existing literature on Tregs in pulmonary fibrosis is inconsistent (42). On one hand, Tregs may contribute to the progression of pulmonary fibrosis by secreting platelet-derived growth factor (PDGF), transforming growth factor-β (TGF-β) and other factors that promote epithelial-mesenchymal transition and alter the Th1/Th2 balance. On the other hand, Tregs may inhibit fibrosis by promoting the repair of epithelial cell damage, inhibiting the accumulation of fibroblasts, and strongly suppressing the production and function of pro-inflammatory factors and cells. The roles of Treg would, thus, be dependent on the time and microenvironment in regards to pulmonary fibrosis. We would like to note that these Tregs express PD-L1. We previously reported the PD-L1 expression in T cells from BALF and the emergence of PD-1 and PD-L1 co-expressing T cells in special situations, such as a severe immune-checkpoint inhibitor-related ILD (43) and adult T cell leukemia-affected lungs (44). These PD-L1 expressing T cells including CD4 T cells may have an immune-suppressing function on neighboring effector T cells or *cis-*regulatory fasion on itself (45) via PD-1/PD-L1 interaction.

We showed increased levels of CXCR3^+^ CD226^+^ CD4^+^ T cells in sarcoidosis. CXCR3 ligands, CXCL9 and CXCL11, are augumented in BALF from pulmonary sarcoidosis (46). These CXCL9 and CXCL11 are interferon-inducible chemokines, and localized to epithelioid histiocytes and multinucleated giant cells forming non-necrotising granulomas (46). Our observation matches the results of the previous study and suggests that CXCR3 ligands may recruite CXCR3-expressing CD4 T cells, which propagate a type 1 immune response and causing granuloma formation. CD226, also known as DNAM-1, is an adhesion molecule that plays a role in activating T cells through the interaction with CD115 and CD112 as ligands (47), competing TIGIT as reciprocal functions. This is the first report to mention CD226 expression in T cells from sarcoidosis. We speculate that CD4 T cells migrating to the lungs through CXCR3 may be activated through CD226, potentially contributing to the pathogenesis of sarcoidosis.

CD57 expression divided T cell subpopulations in viSNE. CD57 expression is typically associated with NK cells, but it has also been found in T cells that have a more advanced differentiation state, reduced replicative capacity, and increased production of cytokines such as IFNγ (48)(49)(50). There is also some evidence that CD57^+^ CD8^+^ T cells may have cytotoxic functions and be correlated with autoimmune activity in type 1 diabetes (51). We found increase levels of CD57^+^CD8^+^PD-1^+^TIGIT^+^ subsets in CTD-ILDs, and these CD8+ subsets may be related to autoimmune condition.

Our study has several limitations, such as the lack of data from healthy controls, the relatively small sample size, and the retrospective design which resulted in missing clinical data for some cases. It is possible that selection biases may be present, as only patients who underwent BAL procedure were able to enroll in this study. In summary, our study demonstrates different immune cell phenotypes in IPF, CTD-ILDs, and sarcoidosis. We discovered that CD14^+^CD36^+^CD84^+^ monocytes and FCRL5^+^ B cells, important immune subsets, are differentially expressed and may be pathogenic. Further studies based on these findings may result in the development of new cell-targeted strategies to inhibit ILD progression.

## Acknowledgements

We thank Ms. Sanae Sekihara PhD. from Standard Biotools for her technical assistance. We also thank the Medical Research Center Initiative for High Depth Omics in Kyushu University.

---

**Supplemental Table 1.** Mass cytometry antibody panels.

For CD45 barcoding
| Label | Target | Product_id | Clone | Amount (uL) |
| --- | --- | --- | --- | --- |
| 89Y | CD45 | 3089003B | HI30 | 1 |
| 106Cd | CD45 | 3106001C | HI30 | 1 |
| 110Cd | CD45 | 3110001C | HI30 | 1 |
| 112Cd | CD45 | 3112001C | HI30 | 1 |
| 116Cd | CD45 | 3116001C | HI30 | 1 |
| 196Pt | CD45 | 3196001C | HI30 | 1 |

For antibody cocktail #1 (T cell panel)
| Label | Target | Product ID | Clone | Amount (uL) |
| --- | --- | --- | --- | --- |
| 141Pr | CD3 | 3141019B | UCHT1 | 1 |
| 142Nd | CD11a | 3142006B | HI111 | 0.5 |
| 143Nd | CD5 | 3143007B | UCHT2 | 1 |
| 145Nd | CD4 | 3145001B | RPA-T4 | 1 |
| 146Nd | CD8a | 3146001B | RPA-T8 | 1 |
| 147Sm | CD7 | 3147006B | CD7-6B7 | 1 |
| 148Nd | CD274 (PD-L1) | 3148017B | 29E.2A3 | 1 |
| 149Sm | CD25 (IL-2R) | 3149010B | 2A3 | 1 |
| 150Nd | CD134 (OX40) | 3150023B | ACT35 | 1 |
| 151Eu | CD2 | 3151003B | TS1/8 | 1 |
| 152Sm | CD95/Fas | 3152017B | DX2 | 1 |
| 153Eu | TIM-3 | 3153008B | F38-2E2 | 1 |
| 154Sm | TIGIT | 3154016B | MBSA43 | 1 |
| 155Gd | CD279 (PD-1) | 3155009B | EH12.2H7 | 1 |
| 156Gd | CD183 (CXCR3) | 3156004B | G025H7 | 1 |
| 158Gd | ST2 (R&D AF523) | Labeling (201158A) |  | 1 |
| 159Tb | CD197 (CCR7) | 3159003A | G043H7 | 1 |
| 160Gd | CD28 | 3160003B | CD28.2 | 1 |
| 161Dy | CD152 (CTLA-4) | 3161004B | 14D3 | 1 |
| 162Dy | CD69 | 3162001B | FN50 | 1 |
| 163Dy | CD272 (BTLA) | 3163009B | MIH26 | 1 |
| 164Dy | CD45RO | 3164007B | UCHL1 | 1 |
| 165Ho | CD223/LAG-3 | 3165037B | 11C3C65 | 1 |
| 166Er | CD44 | 3166001B | BJ18 | 1 |
| 167Er | CD27 | 3167002B | O323 | 1 |
| 168Er | CD278/ICOS | 3168024B | C398.4A | 1 |
| 169Tm | CD19 | 3169011B | HIB19 | 1 |
| 170Er | CD45RA | 3170010B | HI100 | 1 |
| 171Yb | CD226 | 3171013C | DX11 | 1 |
| 172Yb | CD273 (PD-L2) | 3172014B | 24F.10C12 | 1 |
| 173Yb | HLA-DR | 3173005B | L243 | 0.5 |
| 174Yb | CD49d | 3174018B | 9F10 | 1 |
| 175Lu | CD14 | 3175015B | M5E2 | 2 |
| 176Yb | CD57 | 3176019B | HCD57 | 1 |
| 209Bi | CD16 | 3209002B | 3G8 | 1 |

2 For antibody cocktail #2 (B cell/ myeloid cell panel)
| **Label** | **Target** | **Product ID** | **Clone** | **Amount (uL)** |
| --- | --- | --- | --- | --- |
| 141Pr | CD3 | 3141019B | UCHT1 | 1 |
| 142Nd | CD19 | 3142001B | HIB19 | 0.5 |
| 143Nd | CD11b (Biolegend #301302) | Labeling(201143A) | ICRF44 | 0.5 |
| 144Nd | CD38 | 3144014B | HIT2 | 1 |
| 145Nd | CD163 | 3145010B | GHI/61 | 1 |
| 146Nd | CD64 | 3146006B | 10.1 | 1 |
| 147Sm | CD11c | 3147008B | Bu15 | 0.5 |
| 148Nd | IgA | 3148007B | Polyclonal | 1 |
| 149Sm | IgG (Biolegend #410701) | Labeling(201149A) |  | 1 |
| 150Nd | CD138 | 3150012B | DL-101 | 0.5 |
| 151Eu | CD209 (Biolegend #343002) | Labeling(201209A) |  | 1 |
| 152Sm | CD21 | 3152010B | BL13 | 1 |
| 153Eu | CD192 (CCR2) | 3153023B | K036C2 | 1 |
| 154Sm | CD84 | 3154013B | CD84.1.21 | 1 |
| 156Gd | CD86 | 3156008B | IT2.2 | 1 |
| 158Gd | ST2 (R&D AF523) | Labeling (201158A) |  | 1 |
| 159Tb | CD36 (Biolegend #336215) | Labeling(201159A) | 5-271 | 1 |
| 160Gd | CD28 | 3160003B | CD28.2 | 0.5 |
| 164Dy | CD185/CXCR5 | 3164029B | RF8B2 | 1 |
| 165Ho | CD223/LAG-3 | 3165037B | 11C3C65 | 1 |
| 166Er | CD24 | 3166007B | ML5 | 1 |
| 167Er | CD27 | 3167002B | O323 | 1 |
| 168Er | CD206 (MMR) | 3168008B | 44607 | 1 |
| 169Tm | CD32 | 3169020B | FUN-2 | 1 |
| 170Er | TIM-1 (Biolegend #353902) | Labeling(201170A) |  | 1 |
| 171Yb | CD195 (CCR5) | 3171017A | NP-6G4 | 1 |
| 172Yb | IgM | 3172004B | MHM-88 | 1 |
| 173Yb | HLA-DR | 3173005B | L243 | 0.5 |
| 174Yb | IgD (Biolegend #348235) | Labeling(201174A) | IA6-2 | 1 |
| 175Lu | CD14 | 3175015B | M5E2 | 2 |
| 176Yb | antiAPC-176Yb | 3176007B | APC003 | 1 |
| 209Bi | CD16 | 3209002B | 3G8 | 1 |

**Supplementary Figure 1.**
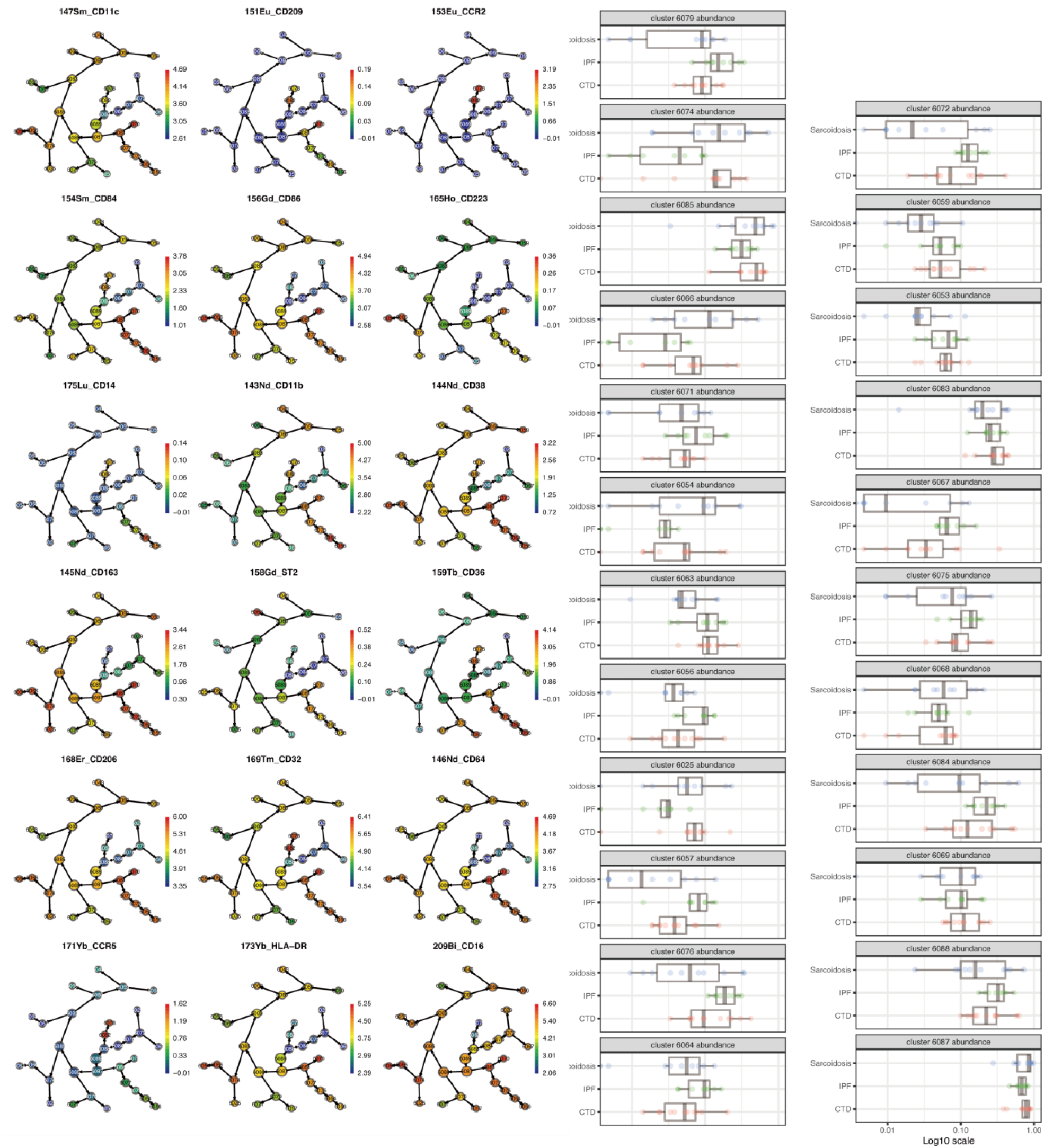
Citrus analysis of myeloid cell populations in BALF cells from IPF, CTD-ILD, and sarcoidosis.

**Supplementary Figure 2.**
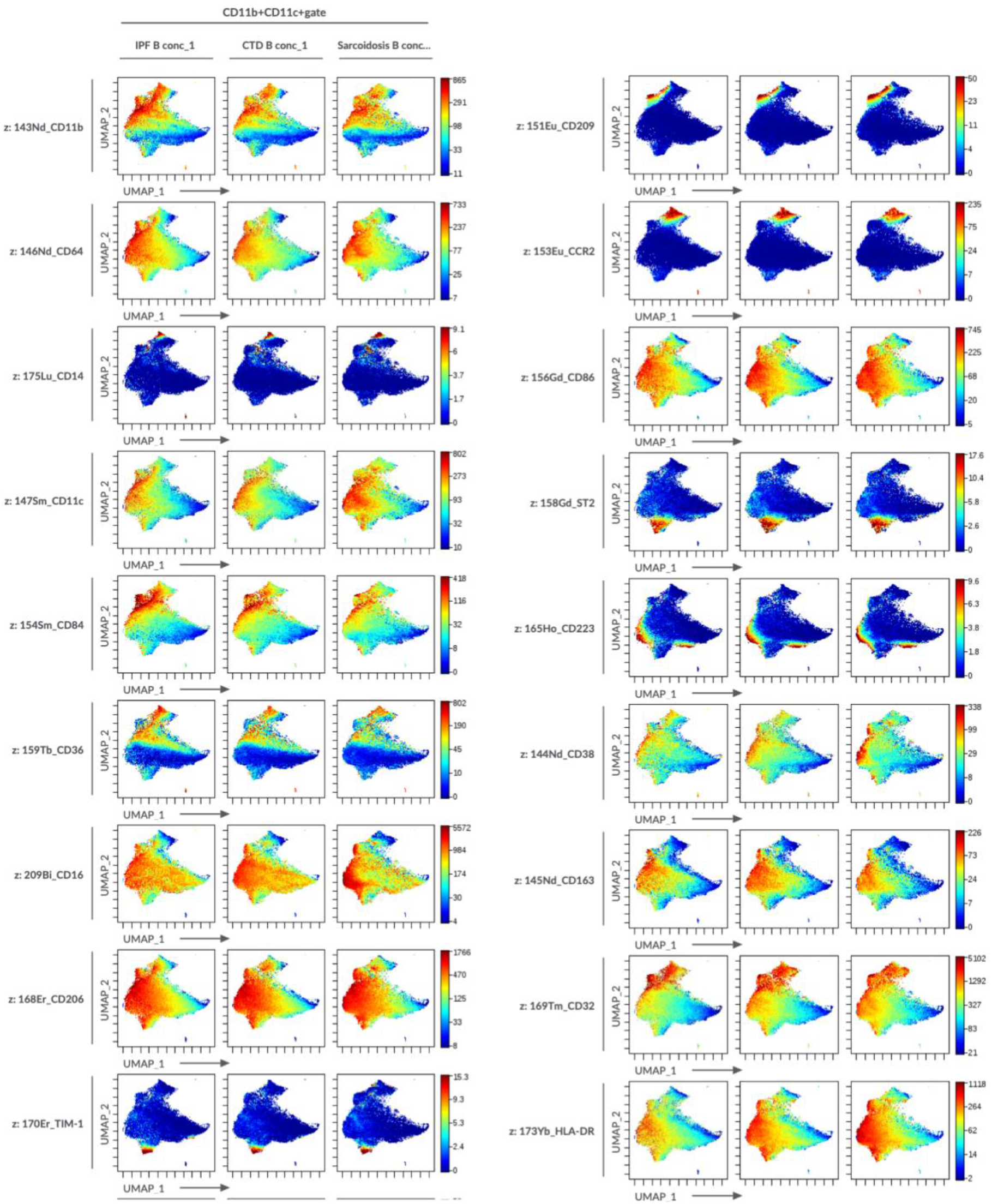
UMAP of concatenated samples visualizing distribution of myeloid cell sub-populations in BALF from patients with IPF, CTD-ILD, sarcoidosis.

**Supplementary Figure 3.**
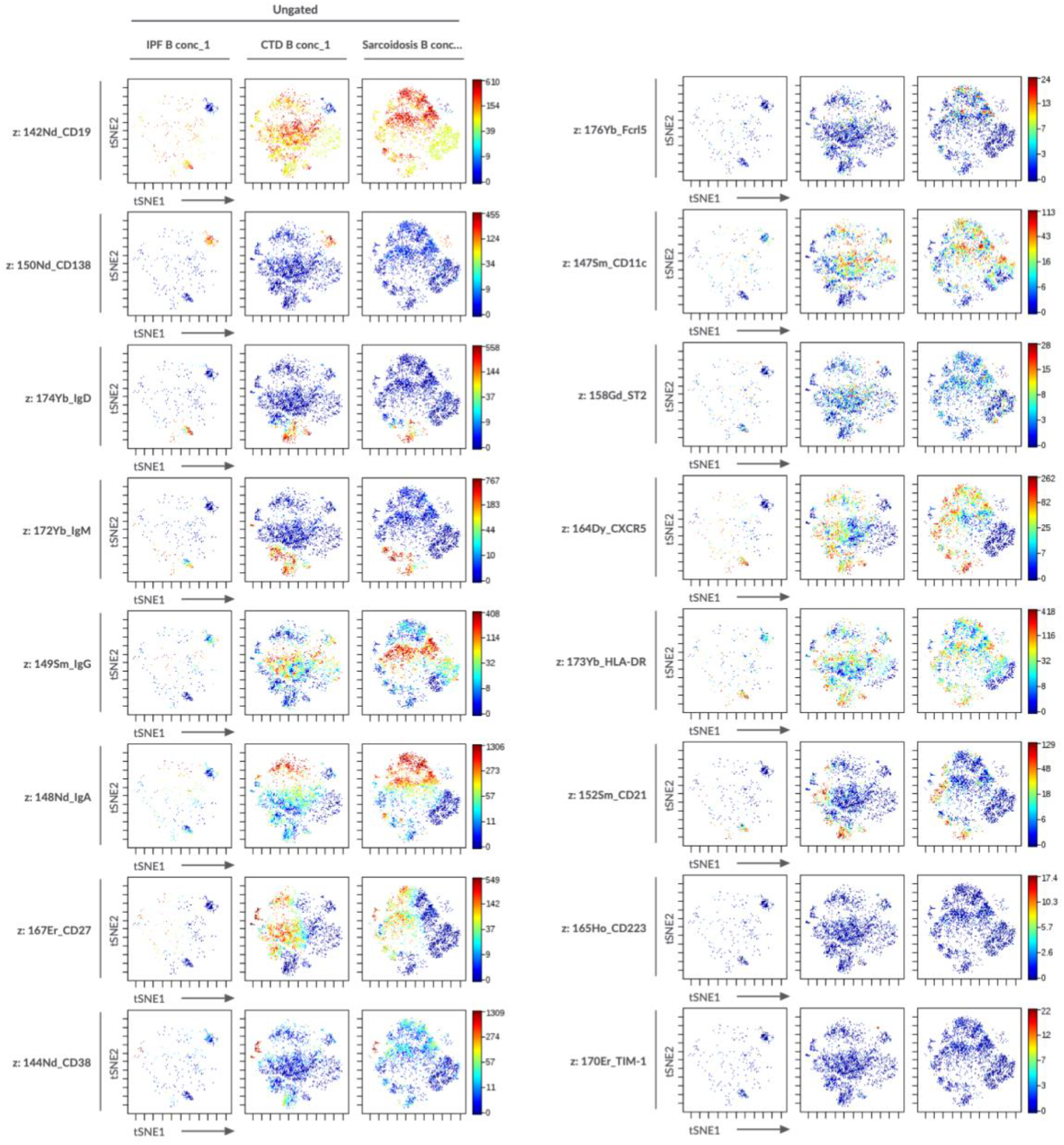
t-stochastic neighborhood embedding (t-SNE) plots of concatenated samples visualizing distribution of B cell sub-populations in BALF from patients with IPF, CTD-ILD, sarcoidosis.

**Supplementary Figure 4.**
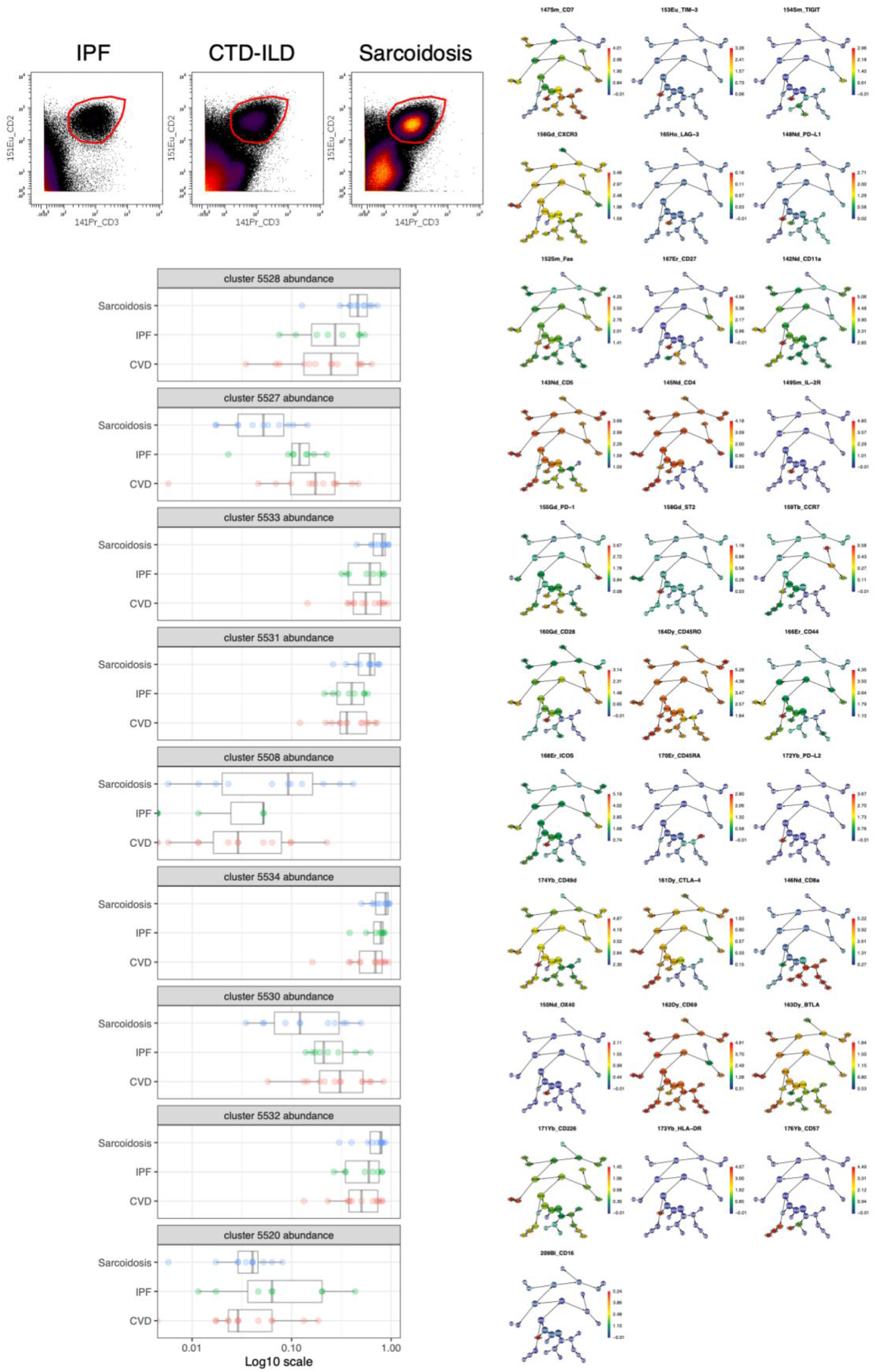
A T cell gate (CD2^+^CD3^+^) and Citrus analysis of T cell populations in BALF cells from IPF, CTD-ILD, and sarcoidosis.

**Supplementary Figure 5.**
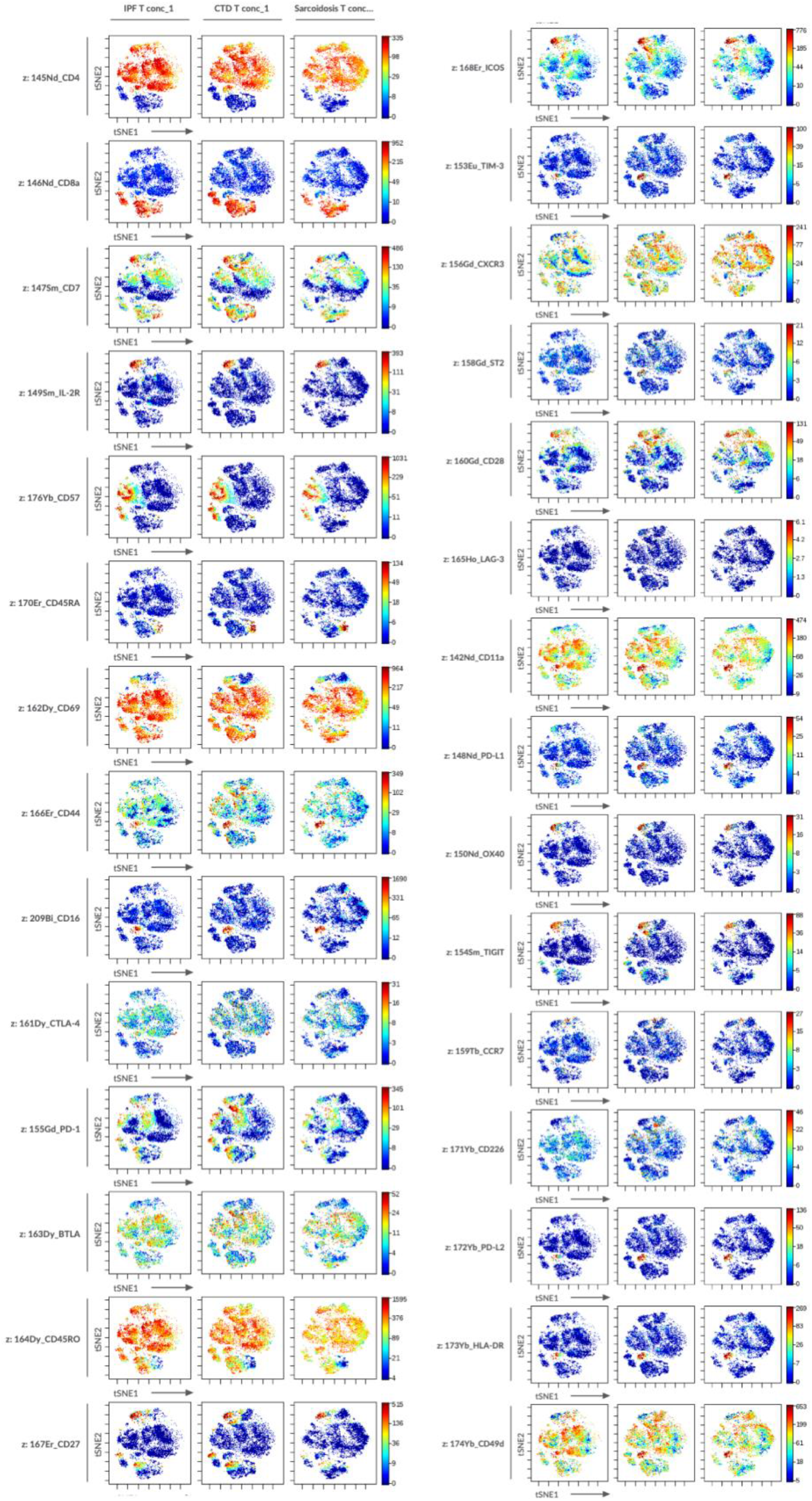
t-stochastic neighborhood embedding (t-SNE) plots of concatenated samples visualizing distribution of T cell sub-populations in BALF from patients with IPF, CTD-ILD, sarcoidosis.

